# KPHMMER: Hidden Markov Model generator for detecting KEGG PATHWAY-specific genes

**DOI:** 10.1101/636290

**Authors:** Hirotaka Suetake, Masaaki Kotera

## Abstract

**Motivation:** Reinforcement of HMMER search for secondary metabolism-specific Pfam domains should contribute to discover novel biosynthetic machinery of clinically important natural products.

**Results:** Here we provide a Python-based command line tool, named as KPHMMER, to extract the Pfam domains that are specific in the user-defined set of pathways in the user-defined set of organisms registered in the KEGG database. KPHMMER outperformed the previous study in detecting secondary metabolism-specific Pfam domain set. Furthermore, it was proven that KPHMMER helps reduce the computational cost compared with the case using the whole Pfam-A HMM file. We believe that KPHMMER is a powerful tool enabling to deal with many other genome-sequenced species for more general purpose.

**Availability:** KPHMMER is implemented as a Python package freely available via the package management system “pip” and also at https://github.com/suecharo/KPHMMER

## 1 Introduction

Metabolic pathways are classified into two categories: *i.e*., those that include ubiquitous chemical compounds such as nucleic acids, proteins and sugars, which are essential for the survival of the living organisms, and those that include species- or clade-specific compounds for the use of interspecies interaction or environmental adaptation, such as toxins or pigments. Even though, the distinction between primary and secondary metabolism is vague (Kotera *et al*., 2008). Some secondary metabolites are important sources of antibiotics, anticancer drugs, *etc*. The genome sequences of various organisms revealed that some actinomycetes have genetic potential to synthesize more secondary metabolites than those known before the genome sequencing (Ohmura *et al*., 2001). This finding indicates that more detailed and comprehensive annotation of secondary metabolism-specific gene sets should allow us to discover novel biosynthetic machinery of clinically important natural products.

Pfam database (Finn *et al*., 2015) contains so-called Pfam domains, which are protein domains conserved structurally and functionally. These Pfam domains are used as clues to annotate functions of proteins or genes. HMMER (Prakash *et al*., 2017) is a widely used tool to search for those domains, and is useful especially for putative proteins that cannot be annotated by BLAST-based homology search (Altschul *et al*., 1997).

The 2ndFind webserver http://biosyn.nih.go.jp/2ndfind/ predicts protein-coding genes using MetaGeneAnnotator (Noguchi *et al*., 2008) or AUGUSTUS (Stanke and Burkhard, 2005), and the obtained putative genes are subjected to HMMER against the Pfam domains of secondary metabolism. In this previous work, 2ndFind prepared gene sets from the genome sequences of four actinomycetes species, and the obtained genes were categorized into primary or secondary metabolism. Pfam domains were categorized into those found in predominantly in primary or secondary metabolism: if a domain was found more frequently in secondary metabolism, then the domain was regarded to be secondary-specific.

**Scheme 1.**
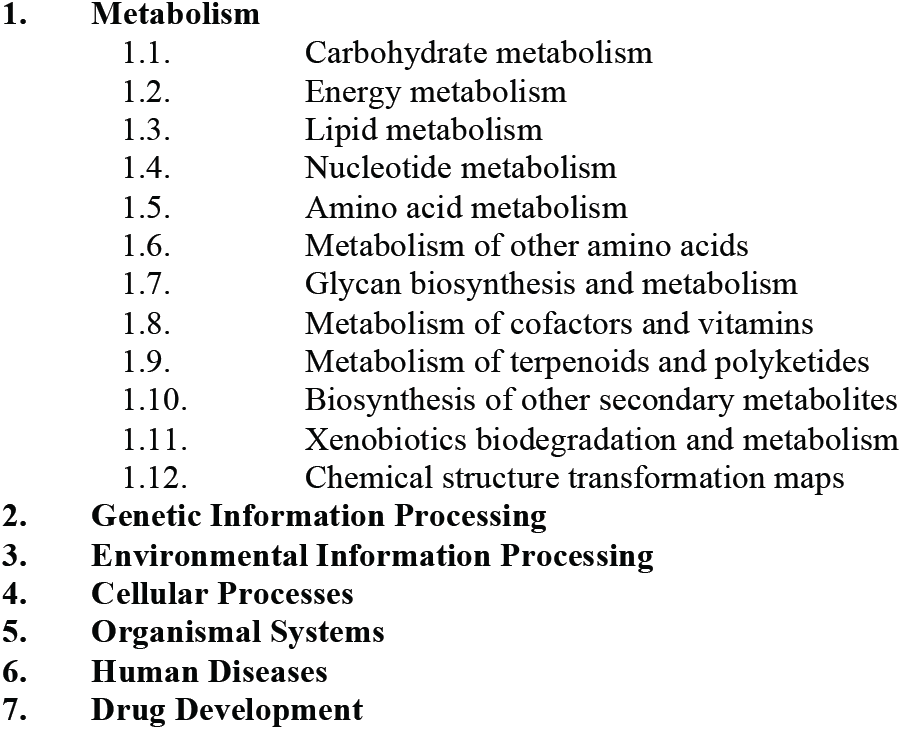
Hierarchy of KEGG PATHWAY

The previous work is valuable for the study of *Streptomyces* and related species, although there still remain rooms to improve. First, the secondary-specific Pfam domains were determined just by comparing the numbers of occurrences in primary and secondary metabolism, even though the fact that the numbers of genes in primary and secondary metabolisms differ substantially. Secondly, the distinction criterion of primary and secondary metabolism was not clear.

Here we provide a Python-based command line tool, named as KPHMMER, to extract the Pfam domains that are specific in the user-defined set of pathways (*e.g*., secondary metabolism) from others (*e.g*., primary metabolism) in the user-defined set of organisms registered in the KEGG PATHWAY database (Kanehisa *et al*., 2016). We show in this paper that KPHMMER outperformed the previous study in detecting secondary-specific Pfam domain set. Furthermore, it was proven that KPHMMER helps reduce the computational cost to extract genes containing the user-defined specific Pfam domain sets in the user-defined set of organisms, compared with the case using the whole Pfam-A HMM file. We believe that KPHMMER is not limited to the study of secondary metabolism, but is a powerful tool enabling to deal with many other genome-sequenced species for more general purpose.

## 2 Methods

### 2.1 Definition of primary and secondary metabolism

We used KEGG PATHWAY database, which classifies pathways into seven classes (Scheme 1). Among them, the first class (1. metabolism) further classifies the pathways into 12 subclasses. In this study, sub-classes 1.9 (Metabolism of terpenoids and polyketides), 1.10 (Biosynthesis of other secondary metabolites) and 1.12 (Chemical structure transformation maps) are regarded as secondary, and the remaining subclasses were regarded as primary. We designed KPHMMER software so that the user can reuse this method to classify pathways into two with the user’s own definitions.

### 2.2 Protein-coding gene sets of six actinomycetes species

The protein-coding gene sets were also obtained from KEGG database, and we designed KPHMMER so that the users can reuse this method for any KEGG-registered organisms of the user’s interest. In this study, for the performance comparison with the previous study, gene sets were retrieved from six actinomycetes species (*Streptomyces avermitilis, Streptomyces coelicolor, Streptomyces griseus, Saccharopolyspora erythraea, Streptomyces scabiei* and *Streptomyces venezuerae*). w

### 2.3 Secondary metabolism-specific Pfam domains and genes

Figure 1 illustrates the proposed procedure to obtain secondary-specific Pfam domains and genes. After categorizing pathways into primary and secondary, the genes in the selected species were classified into primary and secondary gene sets according to the pathways to which the genes belong.

**Figure 1.**
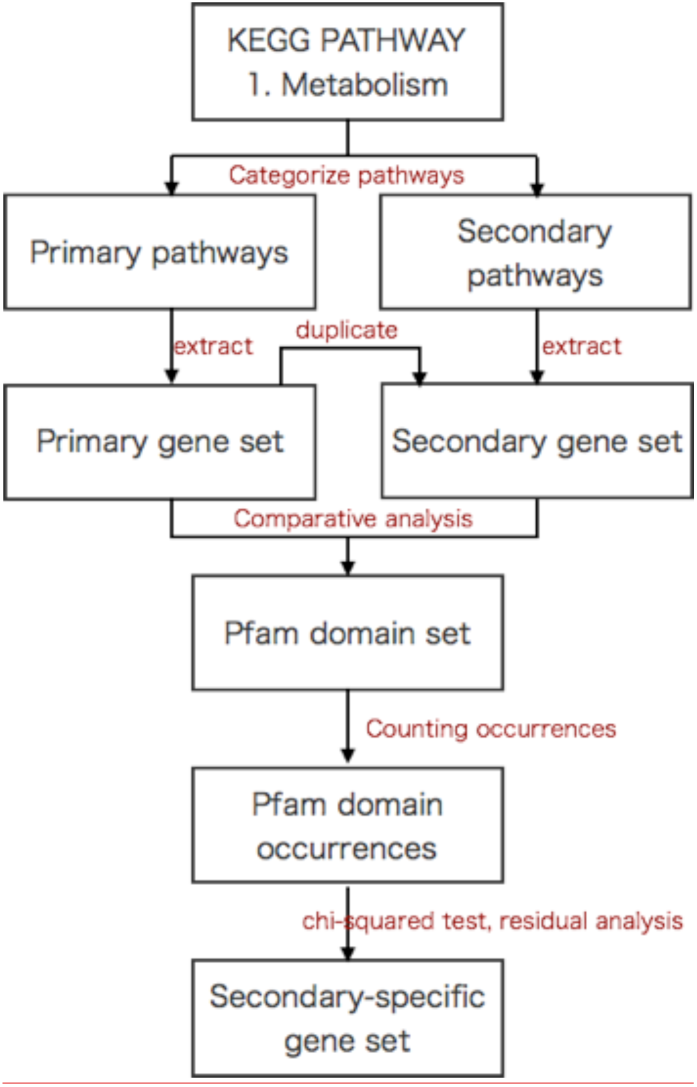
Proposed KPHMMER procedure

In the case where the genes dually appear in primary and secondary pathways, they were classified into secondary (“duplicate” in Figure 1) based on the fact that some “primary” metabolites are used as the intermediate compounds to synthesize secondary metabolites, and some even act as inter-species interaction or environmental adaptation like secondary metabolites. KPHMMER has an option for the user to select which category the user would like to put these dually appeared genes into.

The numbers of occurrences of the genes in primary and secondary gene sets were counted for each Pfam domain. We regarded the Pfam domains and genes as secondary-specific when the occurrence of the Pfam domains was significantly greater in secondary metabolism based on the chi-squared test.

## 3 Results

### 3.1 Implementation of KPHMMER

We implemented a Python-based command line tool named KPHMMER. This tool accepts any genome-sequenced organisms registered in KEGG as the input queries, and outputs the gene set, protein set and Pfam domain set belonging to the category of the user’s interest (*e.g*., the sets significantly found in secondary metabolism). We optimized the API query in this tool to reduce the number of GET request to the KEGG REST API server as many as about 100. This tool also retrieves unstructured KEGG data.

KPHMMER consists of five submethods:

- *search* receives a keyword and search for KEGG organism code (e.g., hsa for *Homo sapiens*).
- *query* receives the KEGG organism code, and outputs the Pfam domains as a yaml file.
- *config* checks or changes the configuration, i.e., the two categories of the pathways to compare, and also the user’s decision which categories the dually appeared genes should be classified into.
- *analysis* receives Pfam domain yaml file, and outputs a tsv file listing Pfam domains or HMM that frequently observed only in the selected category.
- *convert* receives Pfam domain yaml file, and outputs the gene contained therein as fasta file.

An advantage of this tool is its versatility: the user can input any combinations of genome-sequenced organisms (not only the actinomycetes species used in this work), can choose any combinations of pathways of the user’s interest (not only for the distinction of metabolic pathways as shown here, but also of any pathways as listed in Scheme 1), and can obtain Pfam domains or amino acid sequences that specifically occur in the selected set of pathways.

### 3.2 Performance comparison on extracting the secondary-specific domains and genes

According to the previous study, 2ndFind detected 3039 Pfam domains from the 29,728 protein-coding genes found in four actinomycetes species (*Streptomyces avermitilis* ATCC 31267, *Streptomyces coelicolor* A3(2), *Streptomyces griseus* NBRC 13350, *Saccharopolyspora erythraea* NRRL 2338), of which 82 were regarded as secondary-specific Pfam domains (as listed in the webpage http://biosyn.nih.go.jp/2ndfind/), of which three domains (HxxPF_rpt, Lant_dehyd_N, and NRPS) were no longer alive entries but were merged into Condensation (PF00668), Lant_dehydr_N (PF04738), and Condensation (PF00668), respectively. As the result, the number of the current version of secondary-specific Pfam domains was 81.

On the contrary, using the same four actinomycetes species, KPHMMER detected 2,036 Pfam domains from 4,410 metabolism-related genes, of which 28 were regarded as secondary-specific. It was found that the overwrap of the secondary-specific Pfam domains was only five (Aminotran_1_2, KR, NAD_binding_9, ketoacyl-synt, Thiolase_N).

In order to compare the predictive performance of 2ndFind and KPHMMER, we used two taxonomically close *Streptomyces* species, i.e.,*S. scabiei* 87.22 (containing 2,036 Pfam domains in 1,021 metabolism-related genes) and *S. venezuerae* ATCC 10712 (containing 2,036 Pfam domains in 1,068 metabolism-related genes). Both of these *Streptomyces* species are registered in KEGG, and the genes were annotated in either primary or secondary metabolism. This annotation was regarded as the correct data for the performance comparison.

In this study, we used the chi-squared test (p-value < 0.05) to extract secondary-specific Pfam domains in KPHMMER. Table 1 shows the confusion matrices. Performance was evaluated by precision and recall metrics using the numbers of genes classified in primary and secondary metabolisms. Our KPHMMER method outperformed the previous study in both terms of precision and recall in both species. Especially, recall improved dramatically by using KPHMMER. Table 2 shows the top 40 Pfam domains that were regarded as secondary-specific using KPHMMER. This result demonstrates that KPHMMER successfully retrieves some Pfam domains that are known to appear frequently in secondary metabolism but are not listed in 2ndFind (such as KAsynt_C_assoc, and Prenyltransf).

**Table 1.** Performance comparison

|  | TP | FP | FN | TN | Precision | Recall |
| --- | --- | --- | --- | --- | --- | --- |
| *Streptomyces scabiei* |  |  |  |  |  |  |
| 2ndFind | 36 | 122 | 71 | 892 | 0.227 | 0.336 |
| KPHMMER | 100 | 200 | 7 | 814 | 0.333 | 0.934 |
| *Streptomyces venezuelae* |  |  |  |  |  |  |
| 2ndFind | 46 | 114 | 74 | 830 | 0.287 | 0.383 |
| KPHMMER | 113 | 180 | 7 | 764 | 0.385 | 0.941 |

TP (True Positive): the number of genes that belong to secondary metabolism, and were correctly predicted as secondary metabolism. FP (False Positive): the number of genes that belong to primary metabolism, but were falsely predicted as secondary metabolism. FN (False Negative): the number of genes that belong to secondary metabolism, but were falsely predicted as primary metabolism. TN (True Negative): the number of genes that belong to primary metabolism, and were correctly predicted as primary metabolism. Precision: $TP / (TP + FP)$ . Recall: $TP / (TP + FN)$ .

**Table 2.**
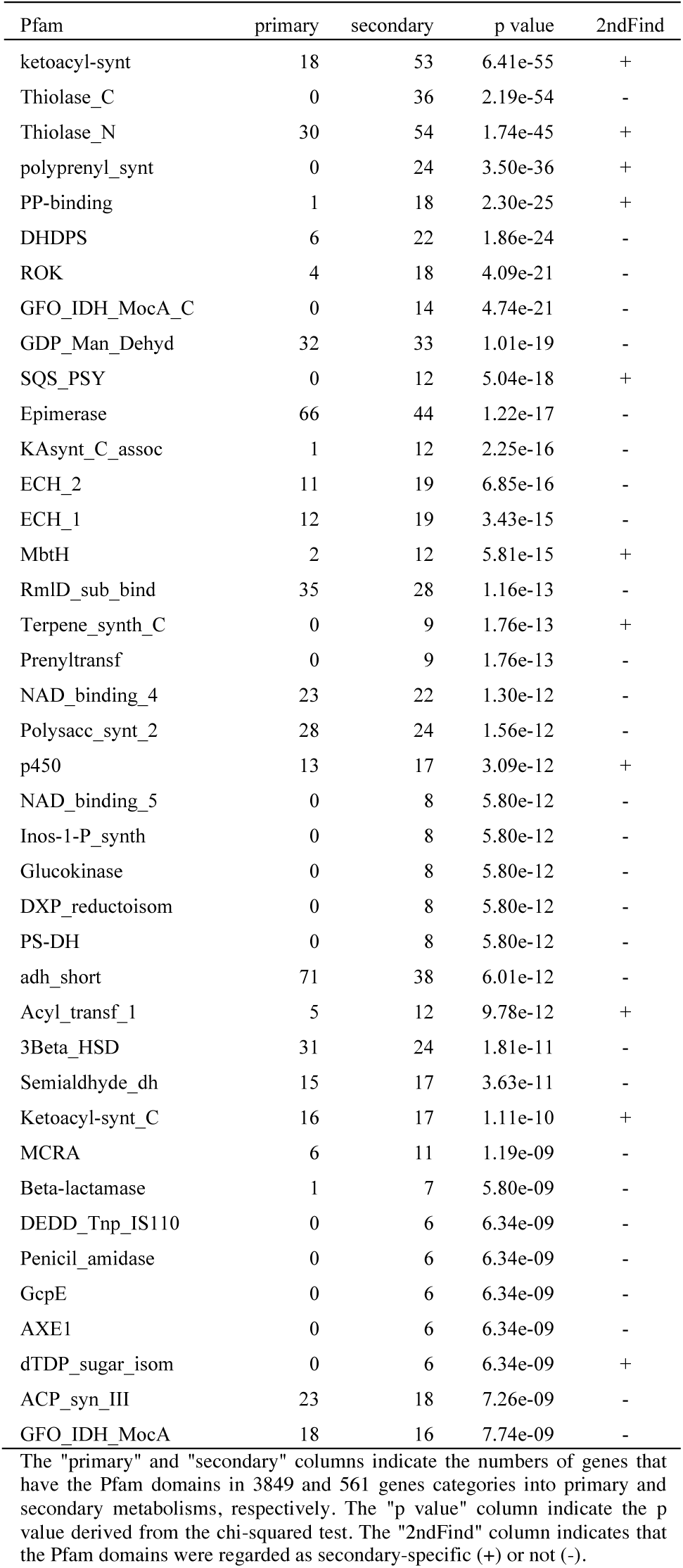
Top 40 Pfam domains that were regarded to be secondary-specific by KPHMMER

### 3.3 Reduction of computational cost by using KPHMMER

Here we further present the effectiveness of our KPHMMER strategy on the reduction of computational costs. Pfam-A.hmm file contains all families of domains defined in Pfam. Pfam 31.0 (as of March 2017, ftp://ftp.ebi.ac.uk/pub/databases/Pfam) contains 16712 entries, and the size of this file is about 1.3 Gb. KPHMMER helps the user reduce the numbers of Pfam domains as the user’s need. For example, when we selected the four actinomycetes species (*Streptomyces avermitilis, Streptomyces coelicolor, Streptomyces griseus, Saccharopolyspora erythraea*) in this study, the obtained HMM file (named as KPHMMER.hmm) only contained 2,036 domains and had 171 Mb.

Table 3 shows the comparison of the computational time to conduct Pfam search. The fasta file used in this study was not very big (∼ 500Kb), and the hmmscan using Pfam-A.hmm took about 11 minutes, compared with about 2 minutes using KPHMMER.hmm. However, this difference should be more serious when the users deal with larger fasta files.

**Table 3.**
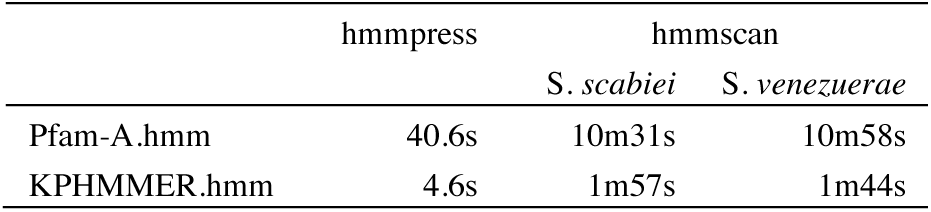
Comparison of the computational time

## 4 Summary

We established a Python-based generic command line tool named KPHMMER, by which the user can generate Pfam domain sets found significantly in the set of pathways of user’s interest, from the organisms of the user’s interest. KPHMMER accepts not only for *Streptomyces* species but also any genome-sequenced species registered in KEGG. The obtained Pfam domain sets should be valuable to the annotation of the genes derived form the closely related species. As the future work, KPHMMER has potential for the enrichment analysis combined with Gene Ontology for transcriptome data, as well as discovering the gene clusters in the genome sequences that contribute to valuable natural product biosynthesis.

## Funding

Japan Society for the Promotion of Science (JSPS) Kakenhi [17K07260].

## Conflict of Interest

none declared.

